## Supplementary Information for "A T6SS in the coral pathogen *Vibrio coralliilyticus* secretes an arsenal of anti-eukaryotic effectors and contributes to virulence"

Supplementary Figures S1-S4

Supplementary Tables S1-S3

Supplementary Datasets S1-S6 captions

Supplementary References

### Supplementary Figures

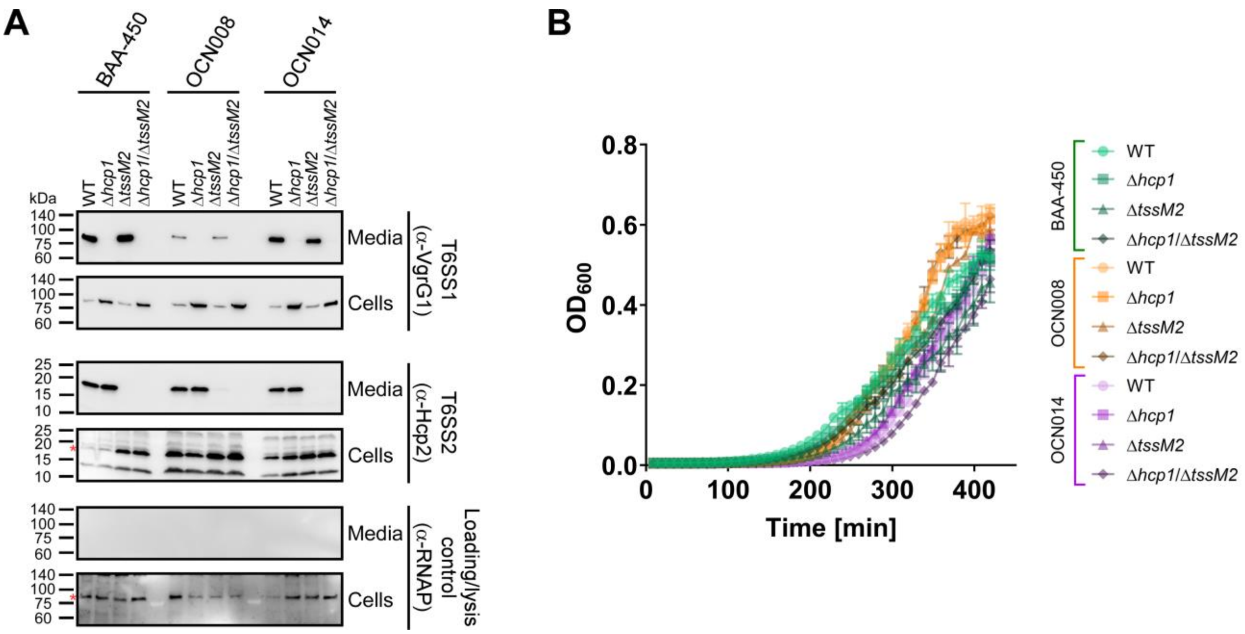


**Fig. S1. Deletion of *hcp1* or *tssM2* inactivates T6SS1 or T6SS2, respectively. A)** Expression (cells) and secretion (media) of VgrG1 and Hcp2 from the indicated *Vcor* strains grown for 4 hours at 28°C in rich media (MLB). RNA polymerase sigma 70 (RNAp) was used as a loading and lysis control. Asterisks denote expected protein sizes. WT, wild-type. **B)** The growth of the indicated *Vcor* strains in MLB at 28°C measured as absorbance at 600 nm ($OD_{600}$). Data are shown as the mean ± SD; $n$ = 3. Results from a representative experiment out of at least three independent experiments are shown.

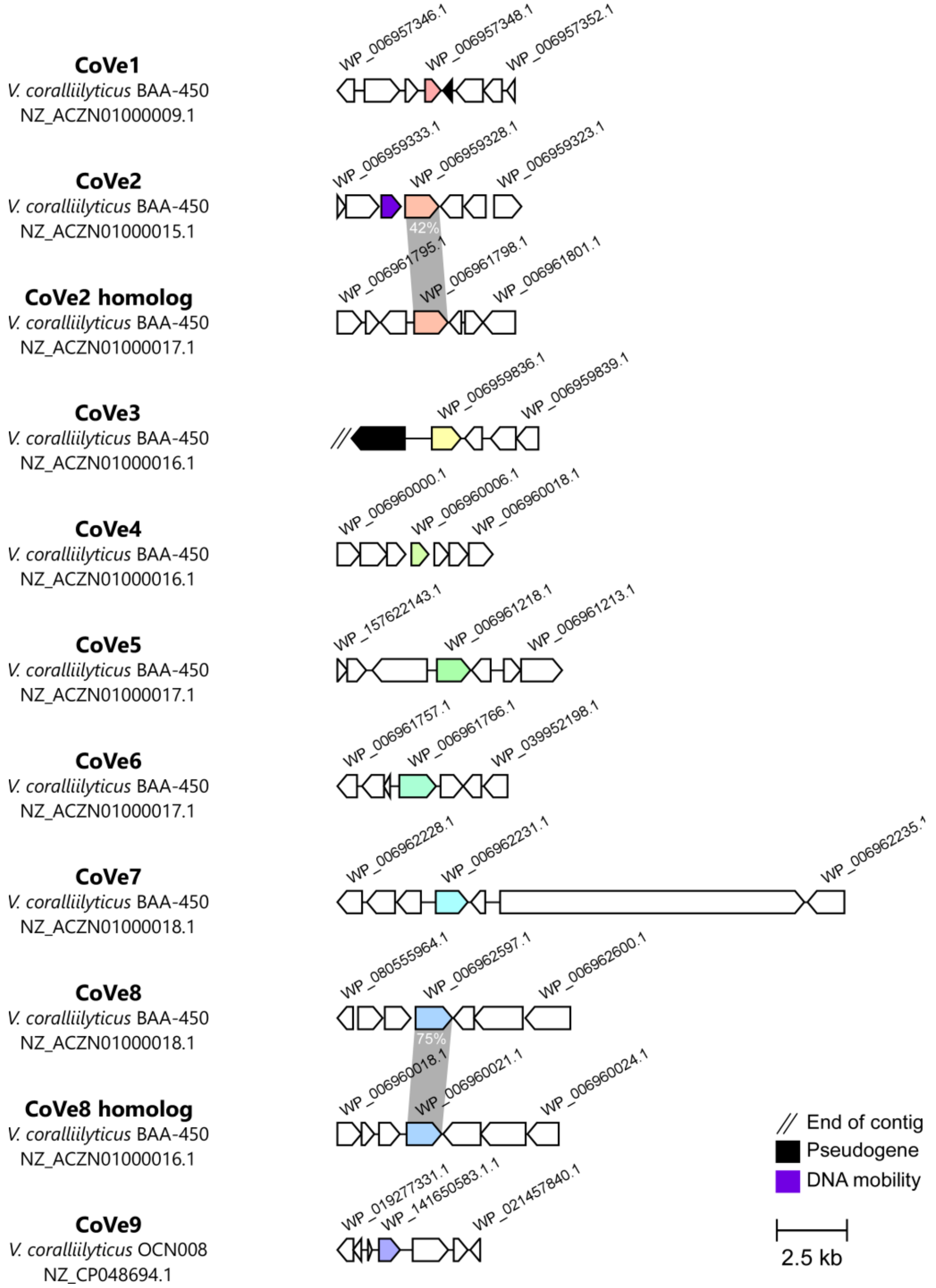


**Fig. S2. Non-structural proteins secreted by *Vibrio coralliilyticus* T6SS2 are encoded by orphan genes.** Genomic neighborhoods of genes encoding representative T6SS effector proteins (CoVes) and their homologs (colored arrows). The strain names, the GenBank accession numbers, and protein accessions are denoted. Genes are denoted by arrows indicating the predicted direction of transcription. Gray rectangles denote regions of amino acid sequence homology; identity percentages are indicated.

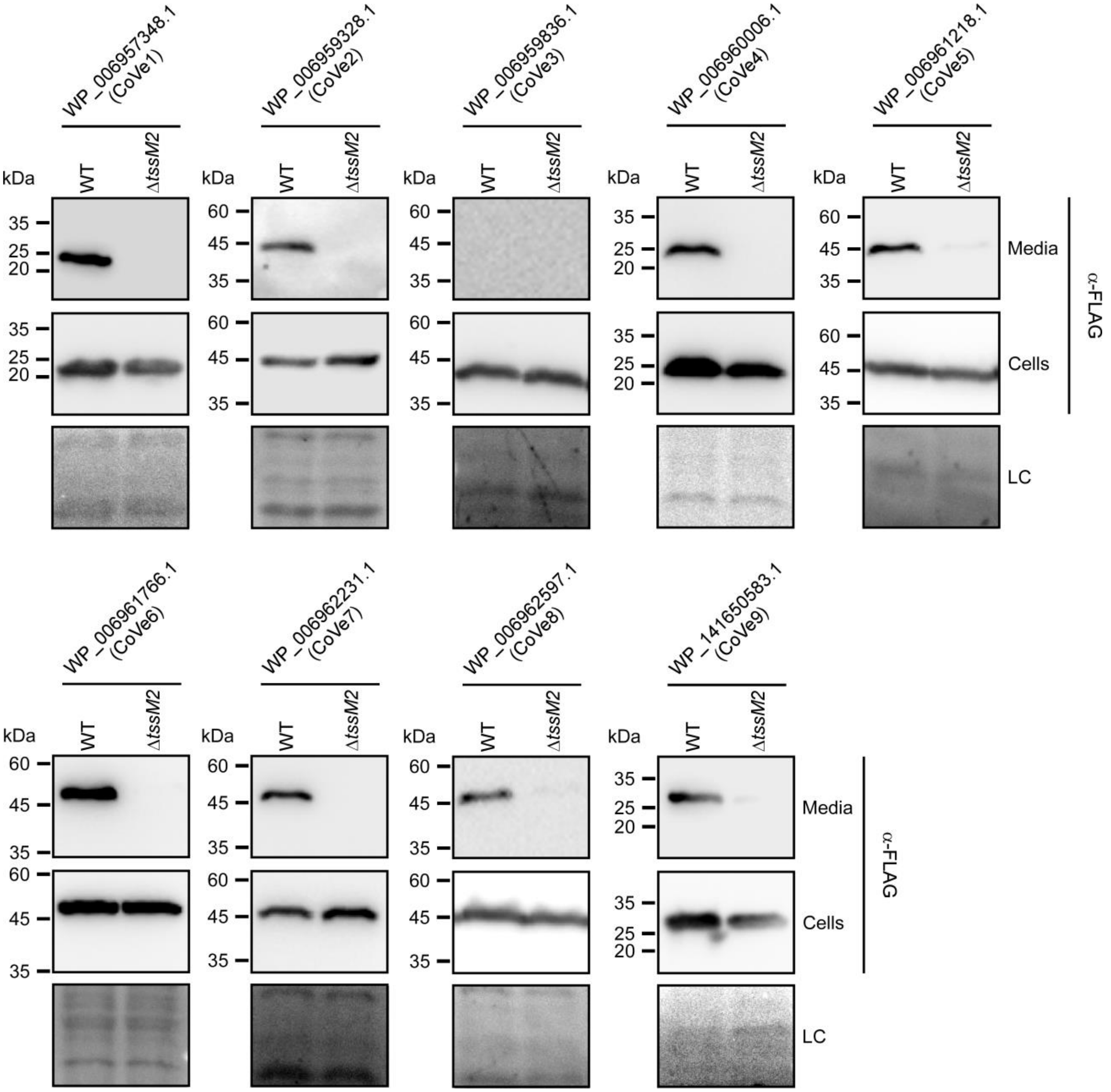


**Fig. S3. CoVes are secreted in a T6SS2-dependent manner.** Expression (cells) and secretion (media) of C-terminally FLAG-tagged CoVes expressed from arabinose-inducible plasmids in *Vcor* strains, either wild-type (WT) or T6SS2⁻ (Δ*tssM2*). CoVe1-8 were monitored in *Vcor* BAA-450 and CoVe9 was monitored in *Vcor* OCN008. *Vcor* strains were grown in MLB supplemented with chloramphenicol and 0.01% (wt/vol) L-arabinose for 4 hours at 28°C. Loading control (LC) is shown for total protein lysate. Results from a representative experiment out of at least two independent experiments are shown.

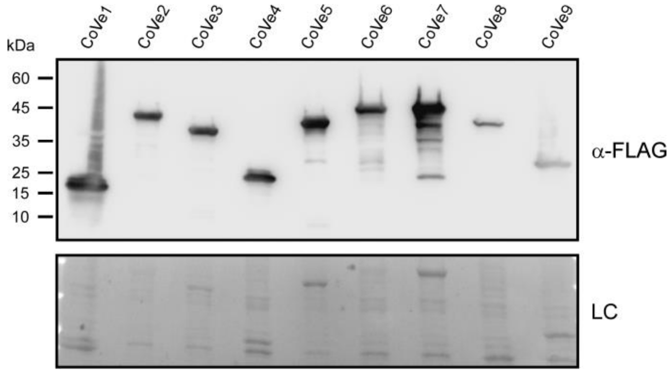


**Fig. S4. CoVes are expressed in *E. coli*.** Expression of C-terminally FLAG-tagged CoVes from arabinose-inducible plasmids in *E. coli* strain DH5α (λ-pir). Loading control (LC) is shown for total protein lysate. Results from a representative experiment out of at least two independent experiments are shown.

### Supplementary Tables

**Table S1. Bacteria and yeast strains used in this study.**

| Strain name | Genotype | Comments | Source |
|---|---|---|---|
| *Vibrio coralliilyticus* BAA-450 | Wild-type | Used for generating deletion strains, as an attacker in competition assays, in BMDM infection assays, and in secretion assays | [1] |
| *Vibrio coralliilyticus* BAA-450 Δ*hcp1* | Δ*VIC_RS16330* | BAA-450 derivative containing a deletion in *hcp1*; used as an attacker in competition assays, in BMDM infection assays, and in secretion assays | [2] |
| *Vibrio coralliilyticus* BAA-450 Δ*tssM2* | Δ*VIC_RS20055* | BAA-450 derivative containing a deletion in *tssM2*; used as an attacker in competition assays, in BMDM infection assays, and in secretion assays | This study |
| *Vibrio coralliilyticus* BAA-450 Δ*hcp1*/Δ*tssM2* | Δ*VIC_RS16330*/Δ*VIC_RS20055* | BAA-450 derivative containing a deletion in *hcp1 and in tssM2*; used as an attacker in competition assays, in BMDM infection assays, and in secretion assays | This study |
| *Vibrio coralliilyticus* OCN008 | Wild-type | Used for generating deletion strains, as an attacker in competition assays, in BMDM infection assays, in *Artemia* infection assays, and in secretion assays | [3] |

| | | | |
|---|---|---|---|
| *Vibrio coralliilyticus* OCN008 Δ*hcp1* | Δ*G3U99_RS23805* | OCN008 derivative containing a deletion in *hcp1*; used as an attacker in competition assays, in BMDM infection assays, in *Artemia* infection assays, and in secretion assays | This study |
| *Vibrio coralliilyticus* OCN008 Δ*tssM2* | Δ*G3U99_13155* | OCN008 derivative containing a deletion in *tssM2*; used as an attacker in competition assays, in BMDM infection assays, in *Artemia* infection assays, and in secretion assays | This study |
| *Vibrio coralliilyticus* OCN008 Δ*hcp1*/Δ*tssM2* | Δ*G3U99_RS23805*/Δ*G3U99_13155* | OCN008 derivative containing a deletion in *hcp1* and in *tssM2*; used as an attacker in competition assays, in BMDM infection assays, in *Artemia* infection assays, and in secretion assays | This study |
| *Vibrio coralliilyticus* OCN014 | Wild-type | Used for generating deletion strains, as an attacker in competition assays, in BMDM infection assays, and in secretion assays | [4] |
| *Vibrio coralliilyticus* OCN014 Δ*hcp1* | Δ*JV59_RS20030* | OCN014 derivative containing a deletion in *hcp1*; used as an attacker in competition assays, in BMDM infection assays, and in secretion assays | This study |
| *Vibrio coralliilyticus* OCN014 Δ*tssM2* | Δ*JV59_31975* | OCN014 derivative containing a deletion in *tssM2*; | This study |

| | | | |
|---|---|---|---|
| | | used as an attacker in competition assays, in BMDM infection assays, and in secretion assays | |
| *Vibrio coralliilyticus* OCN014 Δ*hcp1*/Δ*tssM2* | Δ*JV59_RS20030*/Δ*JV59_31975* | OCN014 derivative containing a deletion in *hcp1 and in tssM2*; used as an attacker in competition assays, in BMDM infection assays, and in secretion assays | This study |
| *Vibrio natriegens* ATCC 14048 | Wild-type | Used as prey in competition assays | ATCC |
| *Escherichia coli* DH5α (λ-pir) | K-12 derivative laboratory strain containing λ-pir | Used for plasmid maintenance, cloning, protein expression, and toxicity assays | Gift from Prof. Eric V. Stabb |
| *Saccharomyces cerevisiae* BY4741 | *MATa his3Δ1 leu2Δ0 met15Δ0 ura3Δ0* | A yeast strain used for protein expression and toxicity assays | Lab stocks |

**Table S2. Plasmids used in this study.**

| Plasmid name | Description | Purpose | Source |
|---|---|---|---|
| pBAD33.1$^F$ | pBAD33.1 with a FLAG tag inserted at the 3' end of the MCS | Used for the arabinose-inducible expression of proteins | [5] |
| psfGFP | pBAD33.1$^F$ plasmid containing the CDS of sfGFP in-frame with the C-terminal FLAG tag | Used to construct the pKara1 plasmid | [6] |
| pWP_006957348.1 | pBAD33.1$^F$ plasmid containing the CDS of WP_006957348.1 from *V. coralliilyticus* BAA-450 in-frame with the C-terminal FLAG tag of the plasmid | Used for the arabinose-inducible expression in *E. coli* and *V. coralliilyticus* | This study |
| pWP_006959328.1 | pBAD33.1$^F$ plasmid containing the CDS of | Used for the arabinose-inducible | This study |

| | | | |
|---|---|---|---|
| | WP_006959328.1 from *V. coralliilyticus* BAA-450 in-frame with the C-terminal FLAG tag of the plasmid | expression in *E. coli* and *V. coralliilyticus* | |
| pWP_006960006.1 | pBAD33.1$^F$ plasmid containing the CDS of WP_006960006.1 from *V. coralliilyticus* BAA-450 in-frame with the C-terminal FLAG tag of the plasmid | Used for the arabinose-inducible expression in *E. coli* and *V. coralliilyticus* | This study |
| pWP_006961766.1 | pBAD33.1$^F$ plasmid containing the CDS of WP_006961766.1 from *V. coralliilyticus* BAA-450 in-frame with the C-terminal FLAG tag of the plasmid | Used for the arabinose-inducible expression in *E. coli* and *V. coralliilyticus* | This study |
| pWP_006962231.1 | pBAD33.1$^F$ plasmid containing the CDS of WP_006962231.1 from *V. coralliilyticus* BAA-450 in-frame with the C-terminal FLAG tag of the plasmid | Used for the arabinose-inducible expression in *E. coli* and *V. coralliilyticus* | This study |
| pWP_141650583.1 | pBAD33.1$^F$ plasmid containing the CDS of WP_141650583.1 from *V. coralliilyticus* OCN008 in-frame with the C-terminal FLAG tag of the plasmid | Used for the arabinose-inducible expression in *E. coli* and *V. coralliilyticus* | This study |
| pWP_006959836.1 | pBAD33.1$^F$ plasmid containing the CDS of WP_006959836.1 from *V. coralliilyticus* OCN008 in-frame with the C-terminal FLAG tag of the plasmid | Used for the arabinose-inducible expression in *E. coli* | This study |
| p WP_006961218.1 | pBAD33.1$^F$ plasmid containing the CDS of WP_006961218.1 from *V. coralliilyticus* OCN008 in-frame with the C-terminal FLAG tag of the plasmid | Used for the arabinose-inducible expression in *E. coli* | This study |
| pWP_006962597.1 | pBAD33.1$^F$ plasmid containing the CDS of WP_006962597.1 from | Used for the arabinose-inducible expression in *E. coli* | This study |

| | | | |
|---|---|---|---|
| | *V. coralliilyticus* OCN008 in-frame with the C-terminal FLAG tag of the plasmid | | |
| pVSV208 | deRed and $Cm^R$ cassette-containing plasmid | Used as the backbone to construct the pKara1 plasmid | [7] |
| pKara1 | pVSV208 containing the araC cassette amplified from psfGFP, upstream of the dsRed gene | Used for the arabinose-inducible expression | This study |
| pKara1:WP_006959836.1 | pKara1 plasmid containing the CDS of WP_006959836.1 from *V. coralliilyticus* BAA-450 replacing sfGFP, in-frame with the C-terminal FLAG tag of the plasmid | Used for the arabinose inducible expression in *V. coralliilyticus* | This study |
| pKara1:WP_006961218.1 | pKara1 plasmid containing the CDS of WP_006961218.1 from *V. coralliilyticus* BAA-450 replacing sfGFP, in-frame with the C-terminal FLAG tag of the plasmid | Used for the arabinose inducible expression in *V. coralliilyticus* | This study |
| pKara1:WP_006962597.1 | pKara1 plasmid containing the CDS of WP_006962597.1 from *V. coralliilyticus* BAA-450 replacing sfGFP, in-frame with the C-terminal FLAG tag of the plasmid | Used for the arabinose inducible expression in *V. coralliilyticus* | This study |
| pGML10 | pGML10 *E. coli*-*S. cerevisiae* shuttle vector, GAL1-10 promoter with a Myc tag at the 3' end of the MCS | Used for galactose-inducible expression in *S. cerevisiae* | Riken |
| pGML10:eGFP | pGML10 containing the CDS of enhanced GFP in the *EcoRI* site of the MCS, in-frame with the C -terminal Myc tag of the plasmid | Used for galactose-inducible expression in *S. cerevisiae* | [2] |
| pGML10:WP_006957348.1 | pGML10 containing the CDS of WP_006957348.1 from *V. coralliilyticus* BAA-450 in-frame with the C- | Used for galactose-inducible expression in *S. cerevisiae* | This study |

| | | | |
|---|---|---|---|
| | terminal Myc tag of the plasmid | | |
| pGML10:WP_006959328.1 | pGML10 containing the CDS of WP_006959328.1 from *V. coralliilyticus* BAA-450 in-frame with the C-terminal Myc tag of the plasmid | Used for galactose-inducible expression in *S. cerevisiae* | This study |
| pGML10:WP_006959836.1 | pGML10 containing the CDS of WP_006959836.1 from *V. coralliilyticus* BAA-450 in-frame with the C-terminal Myc tag of the plasmid | Used for galactose-inducible expression in *S. cerevisiae* | This study |
| pGML10:WP_006960006.1 | pGML10 containing the CDS of WP_006960006.1 from *V. coralliilyticus* BAA-450 in-frame with the C-terminal Myc tag of the plasmid | Used for galactose-inducible expression in *S. cerevisiae* | This study |
| pGML10:WP_006961218.1 | pGML10 containing the CDS of WP_006961218.1 from *V. coralliilyticus* BAA-450 in-frame with the C-terminal Myc tag of the plasmid | Used for galactose-inducible expression in *S. cerevisiae* | This study |
| pGML10:WP_006961766.1 | pGML10 containing the CDS of WP_006961766.1 from *V. coralliilyticus* BAA-450 in-frame with the C-terminal Myc tag of the plasmid | Used for galactose-inducible expression in *S. cerevisiae* | This study |
| pGML10:WP_006962231.1 | pGML10 containing the CDS of WP_006962231.1 from *V. coralliilyticus* BAA-450 in-frame with the C-terminal Myc tag of the plasmid | Used for galactose-inducible expression in *S. cerevisiae* | This study |
| pGML10:WP_006962597.1 | pGML10 containing the CDS of WP_006962597.1 from *V. coralliilyticus* BAA-450 in-frame with the C- | Used for galactose-inducible expression in *S. cerevisiae* | This study |

| | | | |
|---|---|---|---|
| | terminal Myc tag of the plasmid | | |
| pGML10:WP_141650583.1 | pGML10 containing the CDS of WP_141650583.1 from *V. coralliilyticus* BAA-450 in-frame with the C-terminal Myc tag of the plasmid | Used for galactose-inducible expression in *S. cerevisiae* | This study |
| pDM4 | a $Cm^R$ and $ori_{R6K}$-containing suicide vector | Used to generate deletions in *V. coralliilyticus* genomes | [8] |
| pDM4:*tssM2*$^{BAA\text{-}450}$ | pDM4 containing 1 kb downstream and 1 kb upstream of *VIC_RS16330* in its MCS | Used to delete *tssM2* in *V. coralliilyticus* BAA-450 | This study |
| pDM4:*hcp1*$^{OCN008}$ | pDM4 containing 1 kb downstream and 1 kb upstream of *G3U99_RS23805* in its MCS | Used to delete *hcp1* in *V. coralliilyticus* OCN008 | This study |
| pDM4:*tssM2*$^{OCN008}$ | pDM4 containing 1 kb downstream and 1 kb upstream of *G3U99_13155* in its MCS | Used to delete *tssM2* in *V. coralliilyticus* OCN008 | This study |
| pDM4:*hcp1*$^{OCN014}$ | pDM4 containing 1 kb downstream and 1 kb upstream of *JV59_RS20030* in its MCS | Used to delete *hcp1* in *V. coralliilyticus* OCN014 | This study |
| pDM4:*tssM2*$^{OCN014}$ | pDM4 containing 1 kb downstream and 1 kb upstream of *JV59_31975* in its MCS | Used to delete *tssM2* in *V. coralliilyticus* OCN014 | This study |

**Table S3. Primers used in this study.**

| Primer name | Sequence (5' to 3')# | Description |
|---|---|---|
| pDM4_SacI_Gib_R | GAGCTCTCCCGGGATTCCACAAATTG | Used to amplify the pDM4 plasmid backbone for Gibson assembly |
| pDM4_SalI_Gib_F | GTCGACGGTATCGATAAGCTTGATATACAC | |
| OCN008_hcp1_Gib_UP_F | GAATCCCGGGAGAGCTCtgaatttactcag | Used to amplify 1 kb upstream of |

| | | |
|---|---|---|
| OCN008_hcp1_Gib_UP_R | cccttagcgttcggtaagaaaatccttatattccttctaaaaaatttc | *G3U99_RS23805* to construct pDM4:*hcp1*$^{OCN008}$ |
| OCN008_hcp1_Gib_DN_F | gaaatttttagaaggaatataaggattttcttaccgaacgctaaggg | Used to amplify 1 kb downstream of *G3U99_RS23805* to construct pDM4:*hcp1*$^{OCN008}$ |
| OCN008_hcp1_Gib_DN_R | GCTTATCGATACCGTCGACtcaccaatattaaagctg | |
| OCN008/014_hcp1_F_ForPCR | atgccaactcctgcgtatatg | Used to validate deletion of *G3U99_RS23805* from *V. coralliilyticus* OCN008 (with OCN008_hcp1_R_ForPCR) and *JV59_RS20030* from *Vibrio coralliilyticus* OCN014 (with OCN014_hcp1_R_ForPCR) |
| OCN008_hcp1_R_ForPCR | cctttacccattacttcgtg | Used to validate deletion of *G3U99_RS23805* from *V. coralliilyticus* OCN008 (with OCN008/014_hcp1_F_ForPCR) |
| OCN014_hcp1_Gib_UP_F | GAATCCCGGGAGAGCTCtgaatttactcagcg | Used to amplify 1 kb upstream of *JV59_RS20030* to construct pDM4:*hcp1*$^{OCN014}$ |
| OCN014_hcp1_Gib_UP_R | ccttagcgttcggtaagaaaatccttatattccttctaaaaaatttc | |
| OCN014_hcp1_Gib_DN_F | gaaatttttagaaggaatataaggattttcttaccgaacgctaagg | Used to amplify 1 kb downstream of *JV59_RS20030* to construct pDM4:*hcp1*$^{OCN014}$ |
| OCN014_hcp1_Gib_DN_R | GCTTATCGATACCGTCGACtcaccaatattaaagctgtaac | |
| OCN014_hcp1_R_ForPCR | ttcgattgcacggtaagtg | Used to validate deletion of *JV59_RS20030* from *V. coralliilyticus* OCN014 (with OCN008/014_hcp1_F_ForPCR) |
| BAA450_tssM2_Gib_UP_F_(SM) | GAATCCCGGGAGAGCTCtaaattaaagg | Used to amplify 1 kb upstream of *VIC_RS16330* to construct pDM4:*tssM2*$^{BAA\text{-}450}$ |
| BAA450_tssM2_Gib_UP_R_(SM) | cttcagaggctccttaggaatgtgaagataaaccgcgaag | |
| BAA450_tssM2_Gib_DN_F_(SM) | cttcgcggtttatcttcacattcctaaggagcctctgaag | Used to amplify 1 kb downstream of |

| | | |
|---|---|---|
| BAA450_tssM2_Gib_DN_R_(SM) | GCTTATCGATACCGTCGACtgccagaaccagaag | *VIC_RS16330* to construct pDM4:*tssM2*$^{BAA-450}$ |
| OCN008_tssM2_Gib_UP_F_(SM) | GAATCCCGGGAGAGCTCtgccagaacc | Used to amplify 1 kb upstream of *G3U99_13155* to construct pDM4:*tssM2*$^{OCN008}$ |
| OCN008_tssM2_Gib_UP_R_(SM) | cgcggtttatcttcacattcctaaggagcctct | |
| OCN008_tssM2_Gib_DN_F_(SM) | agaggctccttaggaatgtgaagataaaccgcg | Used to amplify 1 kb downstream of *G3U99_13155* from *V. coralliilyticus* OCN008 and *JV59_31975* from *V. coralliilyticus* OCN014 to construct pDM4:*tssM2*$^{OCN008}$ and pDM4:*tssM2*$^{OCN014}$ |
| OCN008/014_tssM2_Gib_DN_R_(SM) | GCTTATCGATACCGTCGACtaaattaaaggcggaag | |
| OCN014_tssM2_Gib_UP_F_(SM) | GAATCCCGGGAGAGCTCaaccagaagatgtcac | Used to amplify 1 kb upstream of *JV59_31975* to construct pDM4:*tssM2*$^{OCN014}$ |
| OCN014_tssM2_Gib_UP_R_(SM) | ggtttatcttcacattcctaaggagcctctga | |
| OCN014_tssM2_Gib_DN_F_(SM) | tcagaggctccttaggaatgtgaagataaacc | Used to amplify 1 kb downstream of *JV59_31975* to construct pDM4:*tssM2*$^{OCN014}$ |
| tssM2_F_ForPCR | ttagatactcgatggtaatgagaaggcag | Used to validate deletion of *VIC_RS16330* from *V. coralliilyticus* BAA-450, *G3U99_13155* from *V. coralliilyticus* OCN008, and *JV59_31975* from *V. coralliilyticus* OCN014 |
| tssM2_R_ForPCR | atggcagaatcaacttcatctaaatctacaaac | |
| pBAD33.1$^{F}$_Gib_ F | GATTACAAGGATGACGACGATAAGTGAAAGCTTGGCTGTTTTGGCGG | Used to amplify pBAD33.1$^{F}$ and pKara1 plasmids backbone with a C-terminal FLAG tag for Gibson assembly |
| pBAD33.1$^{F}$_Gib_ R | CATATGTATATCTCCTTCTTAAAGTTAAACAAAATTATTTCTAGAG | |
| WP_006957348.1_pBAD33.1_F | CTTTAAGAAGGAGATATACATatgcgtccaacgaaagaaaacc | Used to amplify the CDS of WP_006957348.1 to construct pWP_006957348.1 |
| WP_006957348.1_pBAD33.1_R | CGTCGTCATCCTTGTAATCcttaaacgtccttaccgacaacac | |

| | | |
|---|---|---|
| WP_006959328.1_pBAD33.1_F | CTTTAAGAAGGAGATATACATatgactgatacattatcggcgtc | Used to amplify the CDS of WP_006959328.1 to construct pWP_006959328.1 |
| WP_006959328.1_pBAD33.1_R | CGTCGTCATCCTTGTAATCtttggaggcgggactc | |
| WP_006959836.1_pBAD33.1_F | CTTTAAGAAGGAGATATACATatgtataaaaatgatgctattgc | Used to amplify the CDS of WP_006959836.1 to construct pWP_006959836.1 and pKara1:WP_006959836.1 |
| WP_006959836.1_pBAD33.1_R | CGTCGTCATCCTTGTAATCcaagaaatattctaacg | |
| WP_006960006.1_pBAD33.1_F | CTTTAAGAAGGAGATATACATatgaaggacaaaattgttg | Used to amplify the CDS of WP_006960006.1 to construct pWP_006960006.1 |
| WP_006960006.1_pBAD33.1_R | CGTCGTCATCCTTGTAATCggctttagccac | |
| WP_006961218.1_pBAD33.1_F | CTTTAAGAAGGAGATATACATatgtatgaatttttagtacgagac | Used to amplify the CDS of WP_006961218.1 to construct pWP_006961218.1 and pKara1:WP_006961218.1 |
| WP_006961218.1_pBAD33.1_R | CGTCGTCATCCTTGTAATCatctaacactaaacttc | |
| WP_006961766.1_pBAD33.1_F | CTTTAAGAAGGAGATATACATatgattaccgattacgcaaaatatc | Used to amplify the CDS of WP_006961766.1 to construct pWP_006961766.1 |
| WP_006961766.1_pBAD33.1_R | CGTCGTCATCCTTGTAATCataatcctgataggcc | |
| WP_006962231.1_pBAD33.1_F | CTTTAAGAAGGAGATATACATatgaaagtaaaaatcggggttac | |

| | | |
|---|---|---|
| | | Used to amplify the CDS of WP_006962231.1 to construct pWP_006962231.1 |
| WP_006962231.1_pBAD33.1_R | CGTCGTCATCCTTGTAATCgtttggtgactcttc | |
| WP_006962597.1_pBAD33.1_F | CTTTAAGAAGGAGATATACATatgcaagtcaaactacttaaacgtc | Used to amplify the CDS of WP_006962597.1 to construct pWP_006962597.1 and pKara1:WP_006962597.1 |
| WP_006962597.1_pBAD33.1_R | CGTCGTCATCCTTGTAATCcacatccaatggtg | |
| WP_141650583.1_pBAD33.1_F | CTTTAAGAAGGAGATATACATATGtcaaactttctgtttgtaggtg | Used to amplify the CDS of 141650583.1 to construct pWP_141650583.1 |
| WP_141650583.1_pBAD33.1_R | CGTCGTCATCCTTGTAATCtattttcaagcttctg | |
| pGML10_Gib_F | AATTCTGAACAGAAACTGATTTCCGAAGAGGATCTG | Used to amplify the pGML10 plasmid backbone with a C-terminal Myc tag for Gibson assembly |
| pGML10_Gib_R | TCTAGAGTCAATTCGATCCGGGGT | |
| pGML10:WP_006957348.1_F | CGGATCGAATTGACTCTAGAatgcgtccaacgaaagaaaac | Used to amplify the CDS of WP_006957348.1 to construct pGML10:WP_006957348.1 |
| pGML10:WP_006957348.1_R | GGAAATCAGTTTCTGTTCAGAATTcttaaacgtccttaccgacaac | |
| pGML10:WP_006959328.1_F | ACCCCGGATCGAATTGACTCTAGAatgactgatacattatcggcg | Used to amplify the CDS of WP_006959328.1 to construct pGML10:WP_006959328.1 |
| pGML10:WP_006959328.1_R | GGAAATCAGTTTCTGTTCAGAATTtttggaggcgggactcg | |
| pGML10:WP_006959836.1_F | ACCCCGGATCGAATTGACTCTAGAatgtataaaaatgatgctattg | Used to amplify the CDS of WP_006959836.1 to construct pGML10:WP_006959836.1 |
| pGML10:WP_006959836.1_R | GGAAATCAGTTTCTGTTCAGAATTcaagaaatattctaacggcgc | |
| pGML10:WP_006960006.1_F | ACCCCGGATCGAATTGACTCTAGAatgaaggacaaaattgttgc | Used to amplify the CDS of WP_006960006.1 to construct pGML10:WP_006960006.1 |
| pGML10:WP_006960006.1_R | GGAAATCAGTTTCTGTTCAGAATTggctttagccaccgtctttac | |

| | | |
|---|---|---|
| pGML10:WP_006961218.1_F | ACCCCGGATCGAATTGACTCTAGAatgtatgaatttttagtacgag | Used to amplify the CDS of WP_006961218.1 to construct pGML10:WP_006961218.1 |
| pGML10:WP_006961218.1_R | GGAAATCAGTTTCTGTTCAGAATTatctaacactaaacttcgttg | |
| pGML10:WP_006961766.1_F | ACCCCGGATCGAATTGACTCTAGAatgattaccgattacgcaaaatatc | Used to amplify the CDS of WP_006961766.1 to construct pGML10:WP_006961766.1 |
| pGML10:WP_006961766.1_R | GGAAATCAGTTTCTGTTCAGAATTataatcctgataggccccactttc | |
| pGML10:WP_006962231.1_F | ACCCCGGATCGAATTGACTCTAGAatgaaagtaaaaatcggggttac | Used to amplify the CDS of WP_006962231.1 to construct pGML10:WP_006962231.1 |
| pGML10:WP_006962231.1_R | GGAAATCAGTTTCTGTTCAGAATTgtttggtgactcttccagttttaac | |
| pGML10:WP_006962597.1_F | ACCCCGGATCGAATTGACTCTAGAatgcaagtcaaactacttaaac | Used to amplify the CDS of WP_006962597.1 to construct pGML10:WP_006962597.1 |
| pGML10:WP_006962597.1_R | GGAAATCAGTTTCTGTTCAGAATTcacatccaatggtggcatgcc | |
| pGML10:WP_141650583.1_F | ACCCCGGATCGAATTGACTCTAGAatgtcaaactttctgtttgtag | Used to amplify the CDS of WP_141650583.1 to construct pGML10:WP_141650583.1 |
| pGML10:WP_141650583.1_R | TTTCTGTTCAGAATTtattttcaagcttctgggcacctcgtccatc | |

[#] Uppercase, plasmid sequence; lowercase, insert sequence

### Supplementary Dataset captions

**Dataset S1. OrthoANI analysis of *V. coralliilyticus* strains used in this study.**

**Dataset S2. Analysis of *V. coralliilyticus* T6SS gene clusters.**

**Dataset S3. Mass spectrometry results for *V. coralliilyticus* BAA-450 samples.**

**Dataset S4. Mass spectrometry results for *V. coralliilyticus* OCN008 samples.**

**Dataset S5. Mass spectrometry results for *V. coralliilyticus* OCN014 samples.**

**Dataset S6. CoVe distribution in RefSeq *V. coralliilyticus* genomes.**
